# Acetylcholine inhibits platelet activation and regulates hemostasis

**DOI:** 10.1101/324319

**Authors:** John A. Bennett, Sara K. Ture, Rachel A. Schmidt, Michael A. Mastrangelo, Scott J. Cameron, Lara E. Terry, David I. Yule, Craig N. Morrell, Charles J. Lowenstein

**Affiliations:** Aab Cardiovascular Research Institute, Department of Medicine, University of Rochester Medical Center Rochester, NY 14624; Department of Pharmacology and Physiology, University of Rochester Medical Center, 14624

**Keywords:** Alpha-granule, nitric oxide, P-selectin, platelet, thrombosis

## Abstract

Platelets are key mediators of thrombosis. Many agonists of platelet activation are known, but there are fewer identified endogenous inhibitors of platelets, such as prostacyclin and nitric oxide (NO). Acetylcholinesterase inhibitors such as donepezil can cause bleeding in patients, but the underlying mechanisms are not well understood. We hypothesized that acetylcholine is an endogenous inhibitor of platelets.

We measured the effect of acetylcholine or analogues of acetylcholine upon human platelet activation ex vivo. We characterized expression of components of the acetylcholine signaling pathway in human platelets. We tested the effect of a subunit of the acetylcholine receptor, CHRNA7, on acetylcholine signaling in platelets. Acetylcholine and analogues of acetylcholine inhibited platelet activation, as measured by P-selectin translocation and GPIIbIIIA conformational changes. Conversely, we found that antagonists of the acetylcholine receptor such as pancuronium enhance platelet activation. Furthermore, drugs inhibiting acetylcholinesterase such as donepezil also inhibit platelet activation, suggesting that platelets release acetylcholine. We found that NO mediates acetylcholine inhibition of platelets. Human platelets express members of the acetylcholine signaling pathway including *CHRNA2, CHRNA7, CHRNB1*, and *ACHE*. Platelets from mice lacking *Chrna7* are hyperactive when stimulated by thrombin and resistant to inhibition by acetylcholine. Furthermore, acetylcholinesterase inhibitors prolonged bleeding in wild-type mice. Knockout mice lacking *Chrna7* subunits of the acetylcholine receptor display prolonged bleeding as well.

Our data suggest that acetylcholine is an endogenous inhibitor of platelet activation. The cholinergic system may be a novel target for anti-thrombotic therapies.

## Abbreviations

NO: nitric oxide
CHRNA7: cholinergic receptor neuronal nicotinic alpha polypeptide 7
GPIIbIIIA: glycoprotein IIb IIIa
AChR: acetylcholine receptors
AChE: acetylcholinesterase
TRAP: thrombin receptor activating peptide 6
PAR1: protease activated receptor 1
P2Y12: purinergic receptor P2Y
GPVI: glycoprotein VI
NOS3: nitric oxide synthase isoform 3
L-NAME: L-nitroarginine methyl ester

## Introduction

Platelet activation is crucial for hemostasis and thrombosis (1–3). A variety of agonists activate platelets in vivo, including thrombin, collagen, and ADP (4–8). An equally important aspect of platelet biology is inhibition of activation, limiting excess thrombosis which can otherwise lead to stroke or pulmonary embolism. Endogenous platelet inhibitors include factors released from endothelial cells such as nitric oxide and prostacyclin (9–12).

Studies of adverse bleeding reactions to commonly used drugs can reveal novel inhibitors of platelet function (13). For example, a few case reports have suggested that acetylcholinesterase inhibitors are associated with bleeding (14, 15). Several clinical trials have examined the safety of donepezil, and one of these trials showed that donepezil increases the risk of bruising (16, 17). A meta-analysis of clinical trials of acetylcholinesterase inhibitors shows that these drugs increase the risk of bruising by 1.5 fold compared to placebo, although this increased risk is not significant (18). These isolated clinical studies suggest that acetylcholine may be an endogenous inhibitor of platelet activation.

Prior work from other laboratories suggests that acetylcholine receptors (AChR) are involved in platelet function. Human platelets express subunits of the acetylcholine receptor (19). Artificial agonists of AChR stimulate calcium flux across human platelet membranes (19). Agonists of AChR decrease human platelet activation as measured by GPIIbIIIa conformational changes and by aggregation (19). Finally, platelets from mice lacking AChR subunit *Chrna7* have increased activation when stimulated by ADP (20). These important experimental studies suggest that acetylcholine signaling plays a role in inhibiting platelets both in vitro and in vivo.

Gaps remain in our collective knowledge pertaining to the effect of acetylcholine upon platelets. The effect of acetylcholine on platelets stimulated with endogenous agonists other than ADP is not yet completely known. The effect of acetylcholine on platelet degranulation is not fully understood. The effect of endogenous acetylcholine signaling on hemostasis and thrombosis is not well defined. The expression of genes involved in acetylcholine signaling in human platelets is not fully described. And the mechanisms through which clinical drugs targeting acetylcholine affect bleeding in humans has not yet been explored. Determining the role that acetylcholine signaling plays in inhibition of platelet function may help clinicians avoid the toxicity of drugs that target the parasympathetic nervous system, and may help us uncover new pathways which inhibit platelet function.

## Materials and Methods

### Human Platelet Collection

Human blood collection was performed as previously described using protocols approved by the Institutional Review Board at the University of Rochester Medical Center (IRB Protocol RSRB00028659) (21). Normal healthy blood donors were recruited. Subjects were excluded if they had used aspirin or any nonsteroidal anti-inflammatory agent within 10 days before the blood draw. Blood was collected by venipuncture into sodium citrate anticoagulant tubes. Whole blood was centrifuged at 180 × *g* for 15 min to isolate the top layer of platelet-rich plasma (PRP). PRP was diluted 1:20 in room temperature Tyrode’s Buffer (134 mM NaCl, 2.9 mM KCl, 12 mM NaHCO3, 0.34 mM Na2HPO4, 20 mM HEPES, pH 7.0, 5 mM glucose, 0.35% bovine serum albumin) and dispensed in 100 μL volumes for treatment with various drugs.

### Platelet Drug Treatment

Human platelets were suspended in Tyrode’s buffer and placed into microcentrifuge tubes. Drugs were added and the platelets were incubated for 15 min at room temperature. To some samples, L-nitroarginine methyl ester (L-NAME) was added first and incubated for 15 min, then carbachol (Sigma Aldrich) or acetylcholine (Sigma Aldrich) for 15 min, and then TRAP (Tocris Bioscience) or thrombin (Cayman Chemical) for 15 min. Platelets were first treated for 15 minutes with 1,2-bis(o-aminophenoxy)ethane-N,N,N′,N′-tetraacetic acid (BAPTA) and trifluoperazine (TFP) (Sigma Aldritch) for some experiments. For calcium flux experiments with Fura-2 AM, platelet rich plasma was loaded with Fura-2 AM at 5uM for 1 hour at 37 degrees Celsius, and then further prepared as above to yield platelets loaded with Fura-2. HEK293 cells were also loaded as a positive control. Cells were analyzed on a Flexstation 3 (Molecular Devices) for the 340/380 Fura-2 AM ratio.

### Detection of platelet activation by flow cytometry

Phycoerytherin-labeled antibody to CD62P (P-selectin) (Bectin Dickinson) at a dilution of 1:100 was added to platelets following stimulation or drug treatment for 30 min. Platelets were then fixed in 1% formalin. Surface P-selectin was measured by flow cytometry (LSRII, Becton Dickinson). To detect conformational changes in GPIIbIIIa, FITC-fibrinogen (Abcam) was added for 30 minutes, and platelets were analyzed by flow cytometry. We have previously used these techniques to measure platelet activation (22)

### Quantification of cGMP levels by ELISA

Platelets were treated and stimulated as described above. The reactions were stopped and cells lysed by the addition of HCl to a final concentration of 0.1 M. Samples were cleared by centrifugation (14,000 rpm) for 20 minutes. Samples were then analyzed for cGMP content using a commercially available ELISA (Cayman Chemical).

### qPCR for detection of ACh receptor gene expression

qPCR was performed as previously described (23). Murine tissue was harvested immediately following euthanasia and placed on dry ice. Total RNA was isolated with a kit (RNEasy, Qiagen) following the manufacture’s protocol. The A260/A280 ratio of all samples were between 1.9-2.1 as measured by spectrophotometry (NanoDrop; Thermo Scientific). cDNA was synthesized using a kit (RT^2^ Reverse Transcription, SA Biosciences). Quantitative real-time PCR was performed by SYBR Green gene expression assay using Prime PCR Supermix (BioRad) for 40 cycles on an CFX96 thermal cycler (Bio-Rad). Three independent experiments were performed for each tissue, and in each experiment SYBR green quantification was repeated in triplicate for each sample. Prime PCR primer sets were purchased from Bio-Rad. Expression results were calculated by ΔΔCT method and were normalized to the reference genes GAPDH and βActin.

### Mice

All animal experiments were performed under protocols approved by the University of Rochester Medical Center Institutional Animal Care and Use Committee. Mice were obtained from The Jackson Laboratories (Bar Harbor, ME), including C57Bl/6J wild-type mice (C57BL/6J) and *Chrna7* null mice (B6.129S7-Chrna7tm1Bay/J), and bred and maintained in a pathogen free housing room with *ad libitum* access to food and water on a 12/12 light dark cycle.

### Collection and Preparation and Analysis of Mouse Platelets

Collection of blood was performed as previously described (22). Murine blood was obtained by retro-orbital bleeding of anesthetized animals into heparinized Tyrode’s buffer (134 mM NaCl, 2.9 mM KCl, 12 mM NaHCO3, 0.34 mM Na2HPO4, 20 mM HEPES, pH 7.0, 5 mM glucose, 0. 35% bovine serum albumin) in Eppendorf tubes. The blood was then centrifuged to yield platelet rich plasma which was then washed in new tubes containing Tyrode’s buffer with 1% PGE2 to prevent platelet activation. Platelets were then pelleted by 5 min 600 × *g* centrifugation at room temperature and the supernatant was discarded. The pelleted platelets were gently resuspended in Tyrode’s buffer and kept at room temperature for further experiments. For platelet activation, diluted platelet suspension was divided into 100 μl aliquots, stimulated with 10 μl PBS or thrombin for 10 min followed by staining with anti-CD62P-FITC (eBiosciences) or FITC-fibrinogen (eBiosciences) for 15 min, and immediately fixed with 2% formalin. The fluorescence intensity was measured on an LSRII flow cytometer (BD Biosciences). The data were analyzed with FlowJo software (Tree Star Inc.).

### Mouse tail bleeding assay

Hemostasis assays were performed as previously described (22). In brief, mouse tail bleeding time was measured by placing mice on nose cones delivering 2.5% v/v isoflurane. The distal 3 mm of the tail was amputated by a single cut from a sterile razor blade, the tail was immersed immediately in 37°C PBS, and the time to visual cessation of bleeding for 30 sec or continuous bleeding to 20 min maximal duration, whichever occurred first, was recorded.

### Statistical analyses

Data were analyzed by two-tailed Student’s t-test for comparison of two groups, and by Bonferroni corrected two-way ANOVA to compare means of three or more groups. Statistical significance was defined as P < 0.05.

### Study approval

All in vivo procedures on mice were approved by the Division of Laboratory Animal Medicine at the University of Rochester Medical Center. Human blood collection was performed using protocols approved by the Institutional Review Board at the University of Rochester Medical Center.

## Results

### Acetylcholine receptors regulate platelet activation

Since patients taking acetylcholine inhibitors have an increased risk of bleeding, we hypothesized that increased acetylcholine signaling directly inhibits platelet activation. To test this hypothesis, we first analyzed the effect of carbachol, an analog of acetylcholine, on platelet activation. We treated human platelets with increasing concentrations of carbachol, and then stimulated the platelets with the thrombin receptor agonist thrombin receptor activating peptide 6 (TRAP). Carbachol inhibits human platelets activation in a dose dependent manner (Figure 1A). We next explored the effect of acetylcholine on platelet activation. Acetylcholine inhibits TRAP activation of human platelets in a dose responsive manner by over 25% of maximal stimulation (Figure 1B), and acetylcholine inhibits platelet activation over a range of TRAP doses (Figure 1C).

We tested the effect of acetylcholine signaling upon platelets stimulated with different agonists, including: TRAP, which activates the thrombin receptors PAR1 and PAR4; ADP, which activates the ADP receptor P2Y12; U44619 which activates the thromboxane receptor TP; and convulxin, which activates the collagen receptor GPVI. Although carbachol inhibits TRAP activation of human platelets, carbachol has no effect on platelet activation by other agonists (Figure 1D).

The above data show that acetylcholine inhibits alpha-granule release. Next we tested the effect of acetylcholine signaling on other aspects of platelet activation, namely dense granule secretion and GPIIbIIIa conformational changes. We found that the acetylcholine analogue carbachol decreases dense granule exocytosis measured by release of ATP (Figure 1E) and inhibits GPIIbIIIa activation measured by FITC-fibrinogen binding (Figure 1F). Furthermore, endogenous acetylcholine has the same effect (as shown when the acetylcholine esterase inhibitor pyridostigmine is added) (Figure 1E).

Taken together, these data suggest that stimulation of the acetylcholine receptor inhibits platelet activation.

**Figure 1.**
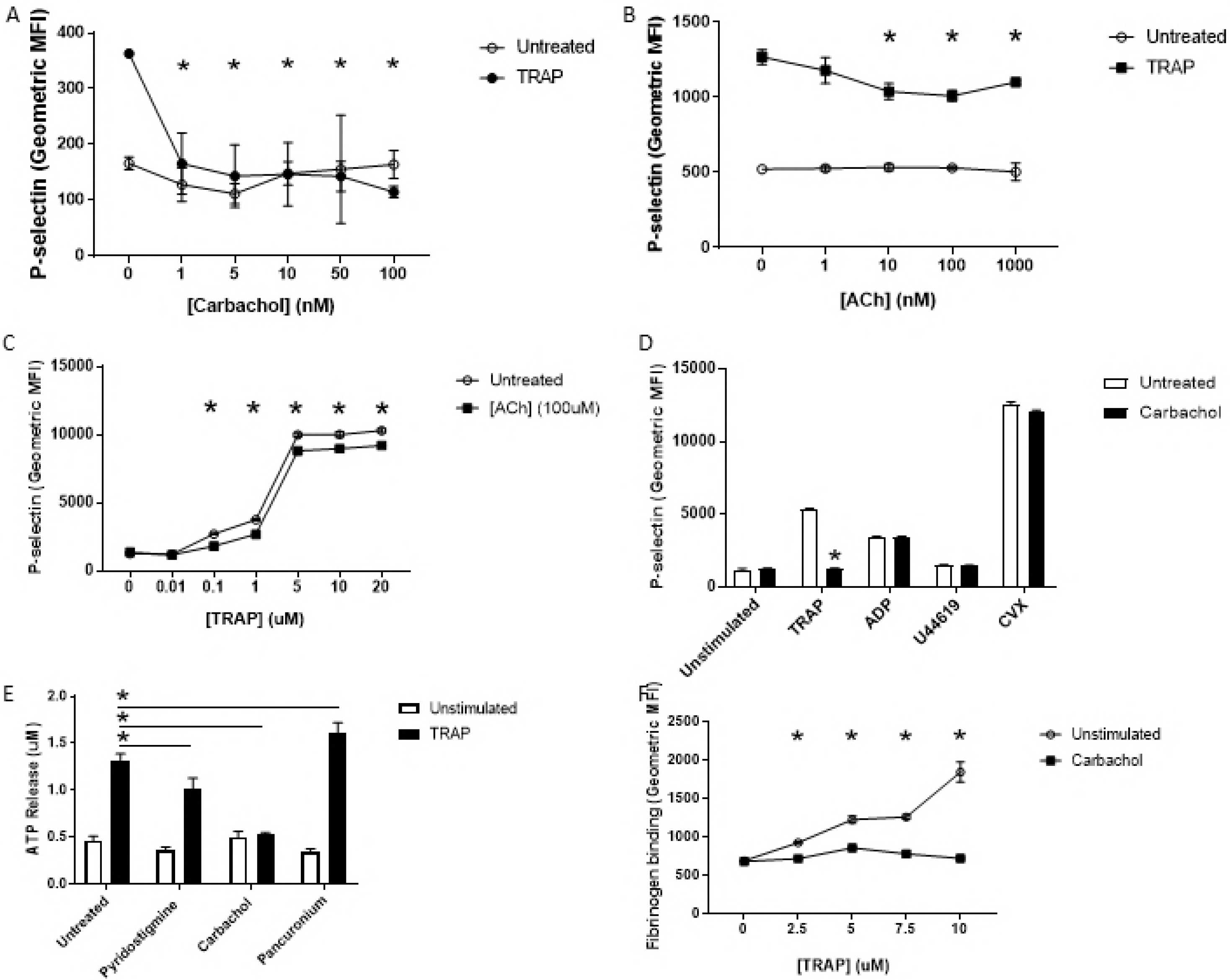
Acetylcholine receptors regulate platelet activation. (A) Carbachol inhibits platelet activation. Human platelets were isolated and treated with PBS or carbachol, stimulated with PBS or 10 uM TRAP, and analyzed for surface expression of P-selectin using flow cytometry. (N=4 ± S.D. *P < 0.05 for TRAP vs. TRAP + carbachol.) (B) Acetylcholine inhibits platelet activation. Human platelets were treated with PBS or ACh, stimulated with PBS or 10 uM TRAP and analyzed as above. (N=4 ± S.D. *P < 0.05 for TRAP vs. TRAP + ACh.) (C) Carbachol inhibits platelet activation over a range of TRAP doses. Platelets were stimulated with varying concentrations of TRAP and analyzed for surface expression of P-selectin as above. (N=4 ± S.D. *P < 0.05 for the indicated concentration of TRAP vs. TRAP + carbachol.) (D) Carbachol inhibits platelet activation by TRAP but not by ADP, U44619, or convulxin. Isolated human platelets were treated with PBS or 10 nM carbachol, then stimulated with various agonists, and analyzed via flow cytometry. (N=4 ± S.D. *P < 0.05 for agonist vs. agonist + carbachol.) (E) Carbachol inhibits platelet dense granule release. Platelets were isolated and treated with 10 nM carbachol, 100 uM pyridostigmine bromide or 100 nM pancuronium bromide, and then stimulated with PBS or TRAP and analyzed for surface expression of P-selectin. (N=4 ± S.D. *P < 0.05 for TRAP vs. TRAP and indicated compound.) (F) Carbachol inhibits GPIIbIIIa activation as measured by FITC-fibrinogen binding to platelets. Platelets were isolated and treated with 10 nM carbachol, and then stimulated with the indicated concentrations of TRAP and analyzed for surface expression of P-selectin. (N=4 ± S.D. *P < 0.05 for TRAP vs. TRAP + carbachol.)

### Endogenous acetylcholine inhibits platelet activation

While acetylcholine signaling inhibits platelet activation, the potential source of acetylcholine in vivo remains unclear. We hypothesized that platelets release acetylcholine which inhibits platelet activation in an autocrine or paracrine manner. We treated platelets with the acetylcholinesterase inhibitor pyridostigmine bromide prior to activation. We observed that inhibition of acetylcholinesterase (AChE) decreases platelet activation (Figure 2A). This is consistent with the idea that pyridostigmine bromide inhibits acetylcholinesterase, increasing the amount of acetylcholine released by platelets which is available to signal through the acetylcholine receptor. We then confirmed that pancuronium bromide, which antagonizes the acetylcholine receptor, enhances platelet activation (Figure 2B). We tested these effect of these compounds on platelet GPIIbIIIa activation using FITC-fibrinogen, and observed that agonism of acetylcholine receptors inhibits, and antagonism of acetylcholine receptors enhances binding (Figure 2C).

Patients who take donepezil may have an increased risk of bleeding (14, 16, 17). Since donepezil is an acetylcholinesterase inhibitor, we hypothesized that donepezil inhibits platelet activation. To test this hypothesis, we treated platelets with donepezil hydrochloride and then stimulated them with TRAP. Donepezil inhibits platelet activation (Figure 2D). These data are consistent with the hypothesis that endogenous acetylcholine released from platelets inhibits platelet activation.

Collectively, these data suggest platelets can release acetylcholine which limits activation, and endogenous acetylcholinesterase limits the extent of endogenous acetylcholine signaling.

**Figure 2.**
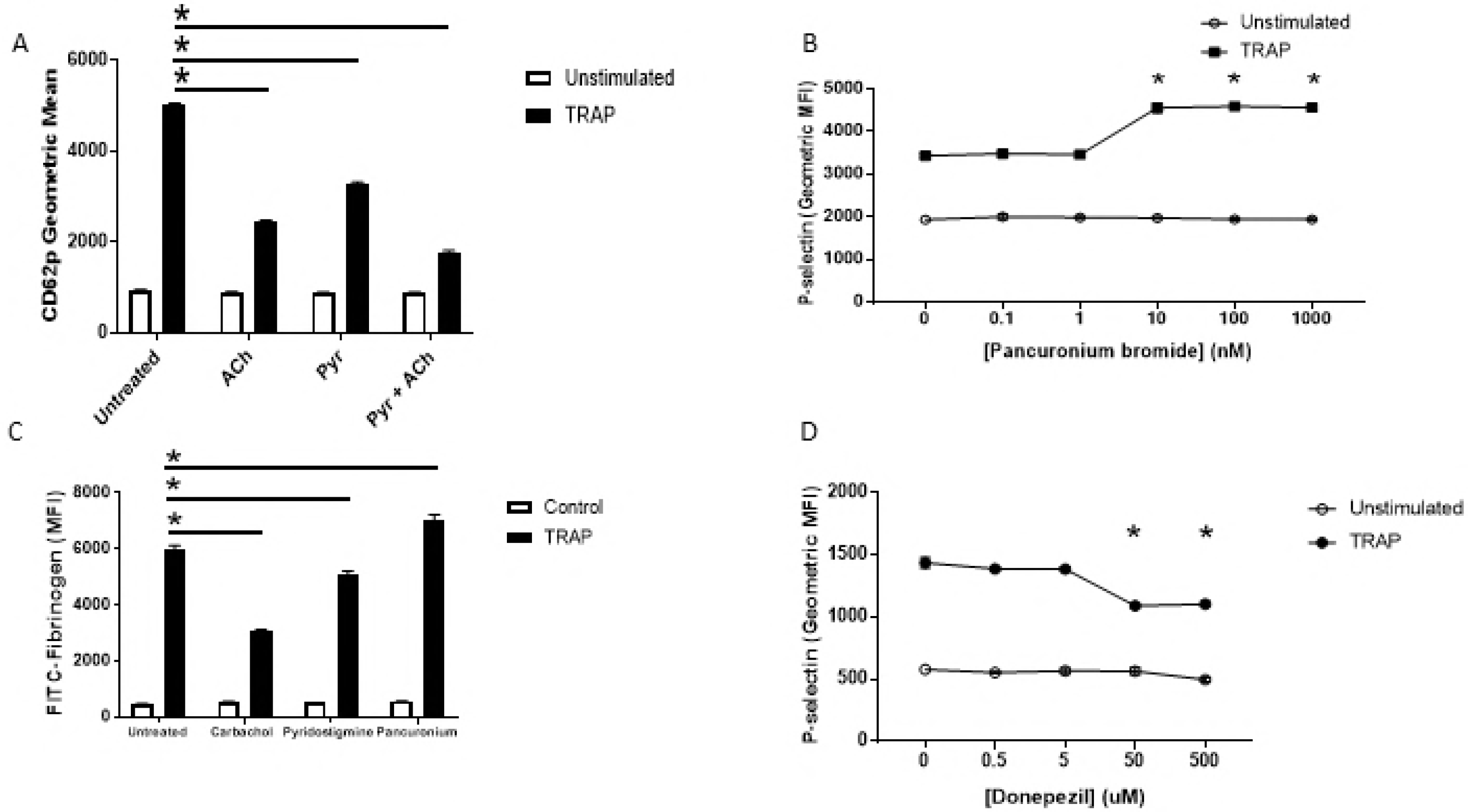
Endogenous acetylcholine inhibits platelet activation. (A) Pyridostigmine inhibition of AChE permits endogenous acetylcholine inhibition of activation of human platelets. Isolated human platelets were treated with 100 uM pyridostigmine, or 100 uM pyridostigmine and 100 uM ACh, stimulated with 10 uM TRAP and then analyzed for P-selectin using flow cytometry. (N=4 ± S.D. *P < 0.05 for TRAP vs. TRAP + pyridostigmine/ACh.) (B) Pancuronium antagonism of acetylcholine receptor blocks endogenous acetylcholine inhibition of human platelets. Isolated human platelets were treated with pancuronium, and then stimulated with 10 uM TRAP and analyzed for P-selectin using flow cytometry. (N=4 ± S.D. *P < 0.05 for TRAP vs. TRAP + pancuronium.) (C) Endogenous ACh inhibits GPIIbIIIa conformational changes. Platelets were isolated and treated with 10 nM carbachol, 100 uM pyridostigmine or 100 nM pancuronium bromide and analyzed for FITC-fibrinogen binding to measure GPIIbIIIa activation. (N=4 ± S.D. *P < 0.05 for TRAP vs. TRAP + indicated compound.) (D) Donepezil inhibition of AChE permits endogenous acetylcholine inhibition of activation of human platelets. Isolated human platelets were treated with donepezil hydrochloride, then stimulated with 10 uM TRAP and analyzed for P-selectin using flow cytometry. (N=4 ± S.D. *P < 0.05 for TRAP vs. TRAP + donepezil.)

### Nitric oxide mediates acetylcholine inhibition of platelet activation

We next explored the mechanism through which acetylcholine signaling inhibits platelet activation. Acetylcholine receptors increase the synthesis of nitric oxide in endothelial cells (24). Platelets express NOS3 (25). We proposed that nitric oxide mediates acetylcholine inhibition of platelets. In order to test our idea, we treated human platelets with an inhibitor of nitric oxide synthase, L-nitroarginine methyl ester (L-NAME), and then treated with carbachol and stimulated with TRAP. We observed that carbachol inhibits platelets, but NOS inhibition blocks the effects of carbachol (Figure 3A). To confirm that acetylcholine signaling triggers NO synthesis in platelets, we measured carbachol stimulation of cGMP, a messenger downstream of NO. Carbachol increases cGMP levels in human platelets, and the effect of carbachol is blocked by the NOS inhibitor L-NAME (Figure 3B). The inhibitory effect of NO was further tested with a range of L-NAME doses. We found that L-NAME inhibits the effects of acetylcholine on platelets in a dose-dependent manner (Figure 3C). Since calcium signaling can regulate NOS activation, we explored a calcium signaling pathway in platelets. First, carbachol increases intracellular calcium levels in platelets (Figure 3D). Second, the calcium chelator BAPTA blocks the ability of carbachol to inhibit platelets. Finally, calmodulin is important for acetylcholine inhibition of platelet activation (Figure 3F). Taken together, our data suggest that NO mediates acetylcholine inhibition of platelets via a calcium-calmodulin dependent mechanism.

**Figure 3.**
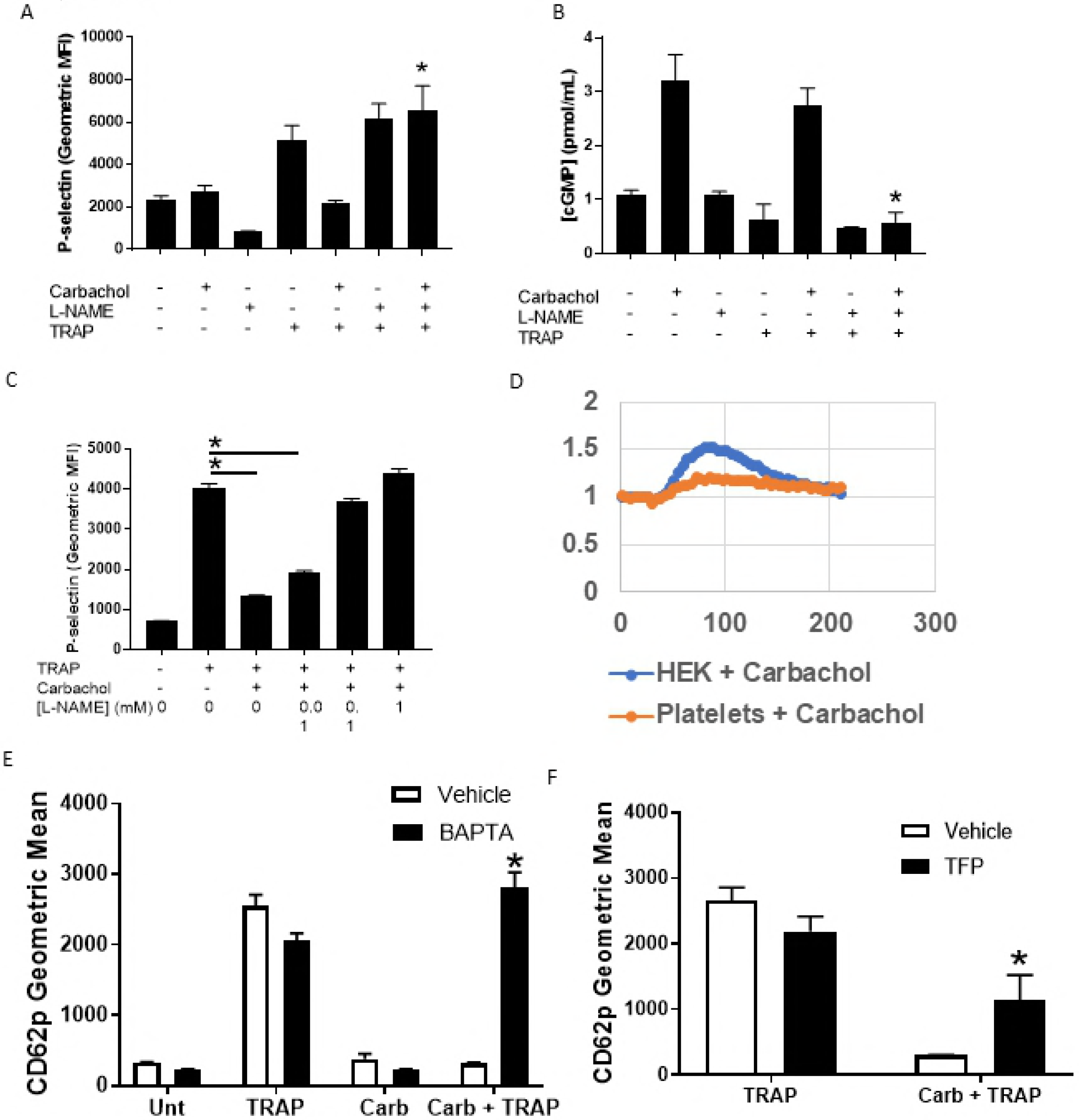
Nitric oxide mediates Ach inhibition of platelet activation. (A) NOS mediates carbachol inhibition of platelet activation. Isolated human platelets were treated with PBS, carbachol, L-NAME or L-NAME + carbachol, stimulated with 10 uM TRAP, and then analyzed for P-selectin using flow cytometry. (N = 4 ± S.D. *P < 0.05 for TRAP + carbachol vs. TRAP + carbachol + L-NAME.) (B) NOS mediates carbachol induced production of cGMP. Isolated human platelets were treated as above, and cGMP content was measured using a commercial kit. (N = 4 ± S.D. *P < 0.05 for TRAP-6 + carbachol vs. TRAP + carbachol + L-NAME.) (C) L-NAME reversal of carbachol mediated platelet inhibition is dose dependent. Platelets were isolated as above and treated with 10 nM carbachol, 1 mM, 0.1 mM or 0.01 mM L-NAME and then stimulated with TRAP and analyzed for surface expression of P-selectin. (N = 4 ± S.D. *P < 0.05 for TRAP + carbachol vs. TRAP + carbachol + indicated concentration of L-NAME.) (D) Carbachol elevates intracellular calcium. Platelets or HEK293 cells were loaded with Fura-2 AM, treated with carbachol and analyzed for calcium flux. (E) Calcium mediates the inhibitory effect of carbachol. Isolated human platelets were treated with BAPTA, then carbachol and then stimulated with TRAP and analyzed for surface expression of p-selectin. (N=4) *P < 0.05 for carbachol + TRAP vs carbachol + TRAP + BAPTA). (F) Calmodulin activity is required for the inhibitory effect of carbachol. Platelets were treated with TFP, then carbachol and then stimulated with TRAP and analyzed for surface expression of p-selectin. *P < 0.05 for TRAP + carbachol vs. TRAP + carbachol + TFP).

### Human platelets express mRNA encoding acetylcholine receptor subunits

We next analyzed human platelets expression of genes involved in acetylcholine signaling and metabolism. Using qPCR of human platelet RNA, we found that human platelets express mRNA for a subunit of the acetylcholine receptor CHRNA7 and acetylcholinesterase ACHE (Figure 4A). Since our qPCR data suggest that human platelets express *CHRNA7* mRNA, we then evaluated if platelets express CHRNA7 protein using a well characterized natural toxin, alpha-bungarotoxin (α-BT), which selectively binds to CHRNA7 subunits. We found that FITC labeled α-BT binds to human platelets, and this binding is specific since the binding can be competitively inhibited by excess non-labeled α-BT (Figure 4B). These data confirm that platelets express functional CHRNA7 subunits on their surface.

**Figure 4.**
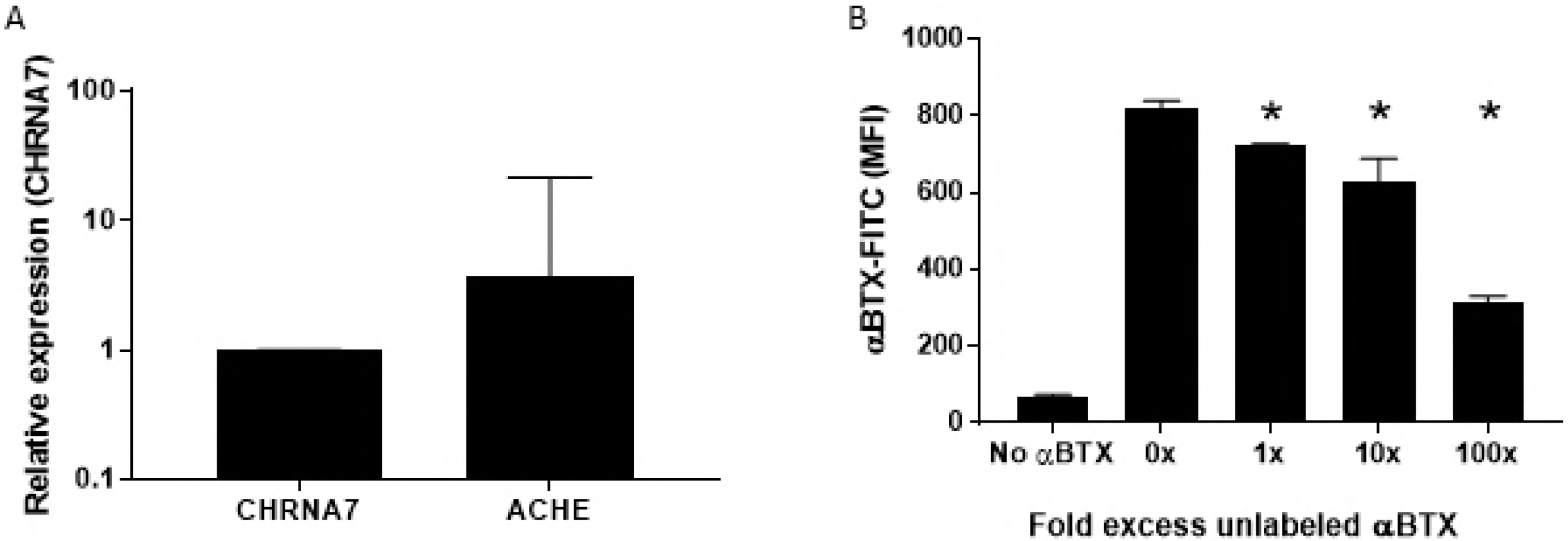
Human platelets express genes related to ACh signaling. (A) Platelets express mRNA transcripts encoding proteins involved in ACh signaling. RNA was isolated from human platelets, a cDNA library was generated, and qPCR was used to evaluate expression of genes related to ACh signaling. Results representative of two separate plate runs, using independent platelet preparations (B) Human platelets express CHRNA7 on their surface. Isolated human platelets were stained with fluorescently labeled αBT in the presence of excess amounts of non-labeled αBT. (N = 4 ± S.D. *P < 0.05 for αBT 0 X vs. 1X or 10 X or 100 X.)

### The acetylcholine receptor subunit CHRNA7 inhibits platelet activation

We next tested the hypothesis that acetylcholine alters platelet activation by signaling through the CHRNA7 subunit of the acetylcholine receptor complex. We treated platelets from *Chrna7* (WT) or *Chrna7* (KO) mice with thrombin and measured their activation by examining P-selectin exposure and fibrinogen binding to the integrin GPIIbIIIA. We observed that *Chrna7*(KO) mice are more sensitive to activation by thrombin indicated by P-selectin exposure (Figure 5A), and that they reach maximum changes in integrin GPIIbIIIa conformation at lower doses of thrombin than WT controls (Figure 5B). This is consistent with the idea that endogenous acetylcholine signaling inhibits platelets.

We also tested the ability of Chrna7 to mediate acetylcholine inhibition of platelet activation. Acetylcholine inhibits activation of platelets from *Chrna7*(WT) mice but fails to inhibit activation of platelets from *Chrna7*(KO) mice (Figure 5C). The acetylcholine receptor agonist carbachol also inhibits activation of platelets from wild-type mice but not from *Chrna7*(KO) mice (Figure 5C).

### Acetylcholine signaling regulates hemostasis in mice

We next tested the effect of acetylcholine signaling in vivo. Clinical case reports suggest that acetylcholinesterase inhibitors are associated with increased risk of bleeding in humans (14, 15). To test the hypothesis that endogenous acetylcholine inhibits hemostasis in vivo, we employed tail transection as a murine model of hemostasis (26). We used two complementary approaches, genetic and pharmacological.

For a genetic approach, we compared *Chrna7*(WT) and *Chrna7*(KO) mice. *Chrna7*(KO) mice have shorter bleeding times than wild-type mice in a tail transection model of hemostasis (Figure 5D).

For a pharmacological approach, we treated mice with donepezil, an AChE inhibitor, and then measured time to cessation of tail bleeding. Donepezil prolongs the bleeding time of mice (Figure 5E). We also confirmed that the acetylcholine receptor agonist carbachol increases bleeding time (Figure 5E).

These data suggest that endogenous acetylcholine inhibits platelet activation and hemostasis in vivo.

**Figure 5.**
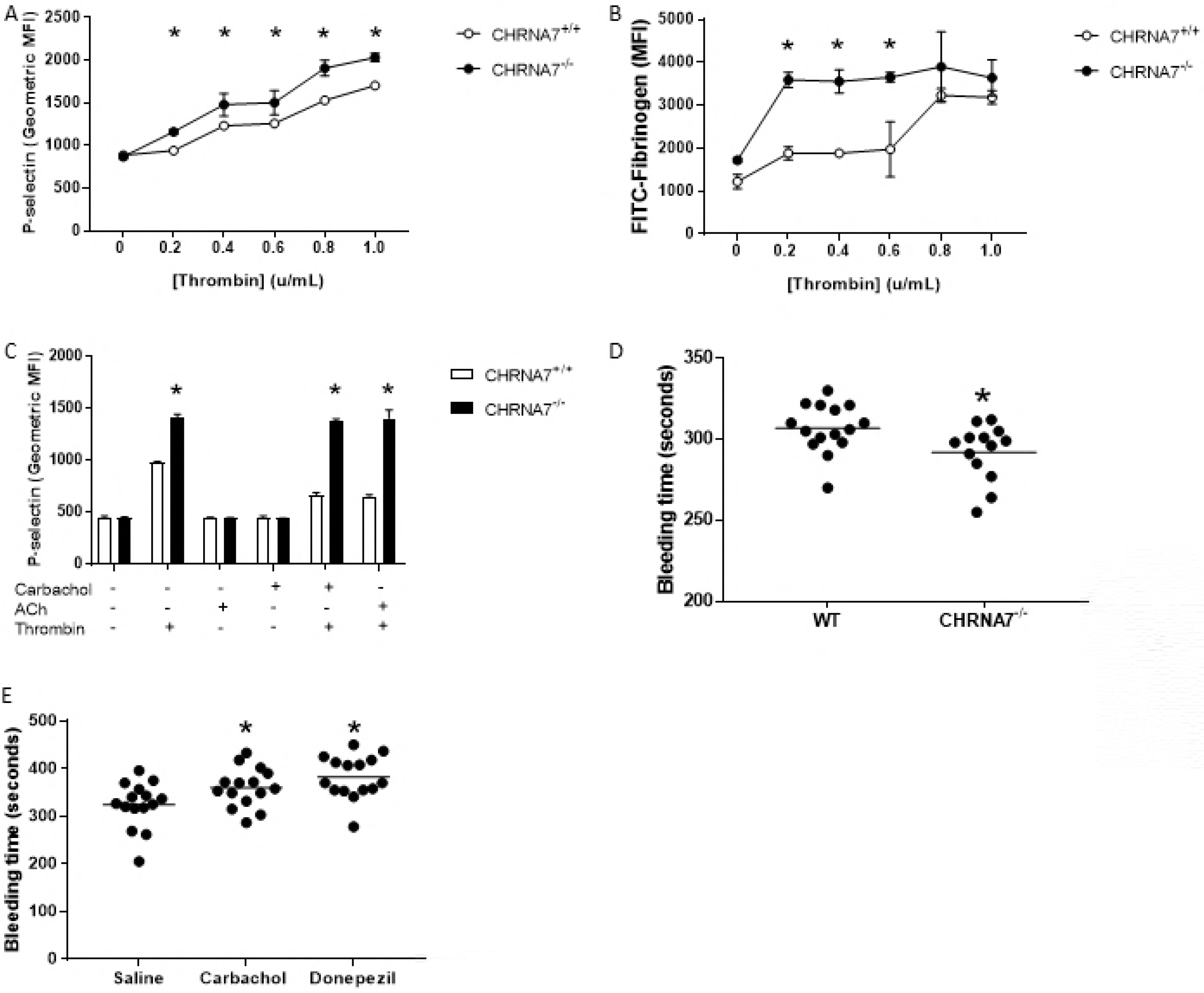
ACh signaling in vivo inhibits hemostasis and platelet activation. (A) Platelets isolated from mice lacking CHRNA7 AChR subunits translocate more P-selectin when stimulated with thrombin. Isolated platelets were treated with thrombin at the indicated concentration and then analyzed for P-selectin using flow cytometry. (N = 4 ± S.D. *P < 0.05 for platelets from *Chrna7*(WT) vs. *Chrna7*(KO) mice.) (B) Platelets isolated from mice lacking CHRNA7 AChR subunits display more active GPIIbIIIa when stimulated with thrombin. Isolated platelets were treated with the indicated concentrations of thrombin, and then analyzed for FITC-fibrinogen binding using flow cytometry. (N = 4 ± S.D. *P < 0.05 for platelets from Chrna7(WT) vs. Chrna7(KO) mice.) (C) Platelets isolated from mice lacking CHRNA7 AChR subunits are not responsive to acetylcholine or carbachol inhibition. Isolated platelets were treated with the indicated concentrations of acetylcholine or carbachol and then stimulated with the indicated concentration of thrombin. (N = 4 ± S.D. *P < 0.05 for platelets from *Chrna7*(WT) vs. Chrna7(KO) mice.) (D) Mice lacking CHRNA7 have prolonged bleeding time. Mice of the indicated genotypes were analyzed for hemostasis using a tail transection assay. (N = 13-15 *P < 0.05 vs. WT control.) (E) Carbachol activation of AChR or donepezil inhibition of AChE prolongs bleeding time in wild-type mice. Mice were treated with 1 mg/kg donepezil or 0.4 mg/kg carbachol and then analyzed for hemostasis using a tail transection assay. (N = 15 *P < 0.05 vs. saline.)

## Discussion

The major finding of our study is that acetylcholine inhibits platelet activation. Acetylcholine signals through the acetylcholine receptor, increasing NO levels, and inhibiting platelet activation. Acetylcholine inhibits activation of platelets from humans and mice by over 15%. Acetylcholine signaling is important in vivo, since mice lacking the acetylcholine receptor subunit *Chrna7* have shorter bleeding times. Finally, an acetylcholinesterase inhibitor drug used in humans that is associated with bruising also inhibits activation of human platelets and prolongs bleeding in mice. Taken together, our results suggest that acetylcholine is an endogenous inhibitor of platelet activation.

We showed that CHRNA7 is necessary for acetylcholine inhibition (Figure 5). Two types of acetylcholine receptors have been described: muscarinic acetylcholine receptors which are G-protein coupled receptors, and nicotinic acetylcholine receptors are ligand gated ion channels (27, 28). Nicotinic acetylcholine receptors are composed of 5 subunits in different combinations, including alpha, beta, delta, epsilon, and gamma subunits (29–31). The precise nature of the acetylcholine receptor in human platelets is not yet defined. Our data suggest that CHRNA7 plays a major role in platelet inhibition by acetylcholine. Further research is needed to identify the subtypes of acetylcholine receptor and their various functions on platelets.

We show that NO mediates acetylcholine inhibition of platelets. Others have demonstrated that platelets express NOS3 and synthesize NO (25, 32, 33). Prior work has shown that NO inhibits platelet adhesion, activation, and aggregation (10, 12, 34–36). For example, we showed that NO inhibits platelet exocytosis (37). Others have shown that activators of NO can inhibit platelet function (38, 39). Our work extends these prior studies and shows that calcium-calmodulin signaling and NOS activity mediate acetylcholine inhibition of platelet activation.

Acetylcholine inhibits activation of platelets by thrombin but not by ADP or thromboxane or convulxin (Figure 4A). There are several possible explanations for this difference. Although both PAR1 and P2Y12 are GPCR, they signal through different intracellular messenger pathways (4, 40–42). Convulxin signals through GPIV (43, 44). While these pathways ultimately converge to stimulate platelet activation as measured by conformational changes in GPIIbIIIa, the prior signaling events are different, and might be differentially susceptible to NO. There are clinical drugs which take advantage of pathway specificity for platelet activation. For example, ticagrelor inhibits platelet activation by thromboxane-A2, but does not inhibit their activation with ADP or collagen (45–47).

We found that acetylcholine inhibits platelet activation in vitro by about 15% (Figure 1B). Carbachol, an analog of acetylcholine, has a much stronger effect upon platelet activation, inhibiting P-selectin translocation by over 90% (Figure 1A and 5A). Thus exogenous agonists like carbachol have a powerful effect upon platelet activation, but endogenous agonists such as acetylcholine have a more modest inhibitory effect on platelet activation.

Our work extends prior research on cholinergic signaling in platelets. Others have shown that agonists of AChR decrease human platelet activation ex vivo as measured by GPIIbIIIa conformational changes and by aggregation (19). We show that acetylcholine itself inhibits platelet degranulation. (Figure 1B). Others have shown that platelets from mice lacking *Chrna7* have increased aggregation when stimulated by ADP ex vivo (20). We extend these data, showing that these platelets do not respond to acetylcholine and have increased exocytosis (Figure 5C). Finally, we add to the current literature by showing that acetylcholine and its analogs modulate hemostasis in vivo.

Our study has several limitations which suggest future studies. We have not yet defined the composition of the acetylcholine receptor on platelets, and we have not identified the role of all acetylcholine subunits in mediating platelet inhibition. Another limitation is that we have indirect evidence that platelets store acetylcholine in their granules, since acetylcholinesterase inhibitors boost platelet inhibition, but we have not directly measured acetylcholine inside platelet granules.

Our studies have pharmacological relevance to humans. We show that donepezil inhibits platelet activation ex vivo at a concentration between 5 - 50 uM (Figure 2D). This matches the concentration of donepezil of 47 uM in serum of humans taking donepezil as a treatment for Alzheimer’s Disease(48). Reports in the literature suggest that drugs targeting the acetylcholine signaling pathway have modest effects on hemostasis; for example, donepezil increase bruising by about 2% more than placebo (18). Our data support our proposal that drugs that target acetylcholine esterase can promote bleeding in humans, and may explain why donepezil is associated with hemostatic abnormalities in humans.

Our study also has therapeutic implications for the management of thrombosis. Our data suggest that drugs targeting acetylcholine receptor subunits might inhibit thrombosis. Furthermore, our data suggest that drugs increasing acetylcholine signaling will increase the risk of bleeding and bruising in patients.

## Acknowledgments

a) Acknowledgements: J. A. Bennett, C. N. Morrell, S. J. Cameron, and C. J. Lowenstein designed the experiments. J. A. Bennett and R. A. Schmidt performed the in vitro analyses of platelets. J. A. Bennett, S. Ture and M. A. Mastrangelo performed the in vivo studies with mice. J. A. Bennett and C. J. Lowenstein wrote the manuscript. C. N. Morrell, S. J. Cameron, and C. J. Lowenstein revised the manuscript. C. J. Lowenstein supervised the research.

b) Sources of Funding: This work supported by grants: R01 HL134894 (CJL), R01 HL124018 (CNM), K08 HL128856 (SJC).

c) Disclosures: The authors declare that no conflicts of interest exist.

